# Sex differences in social synchronization of conditioned fear

**DOI:** 10.1101/2022.10.10.511569

**Authors:** Wataru Ito, Alexei Morozov

## Abstract

Socially coordinated threat responses support the survival of social groups, and the distinct social roles of males and females predict sex differences in such coordination. However, this study area remains unexplored in mice, the most commonly used laboratory species. Here, we examined two behaviors: a recently reported ‘fear synchrony,’ where paired mice synchronize auditory-conditioned freezing more strongly in males, and a newly identified ‘CS-induced affiliation,’ where mice increase proximity upon a conditioned stimulus. These behaviors necessitate the integration of social cues with emotional CS. To understand how sex influences that process, we manipulated social cues through partner familiarity and emotional states via prior stress. Unfamiliarity moderately reduced synchrony in male dyads but not in females. Whereas stress disrupted male synchrony and contrarily enhanced female synchrony. Unfamiliarity eliminated CS-induced affiliation in both sexes, while stress caused males to distance each other but had no effects in females. Interestingly, heterosexual dyads showed resilience in both coordinated behaviors unaffected by stress or unfamiliarity. These findings reveal sex-specific adaptations in socio-emotional integration when orchestrating socially coordinated behaviors and suggest that the sex-recognition circuits confer stress- and unfamiliarity-resilience, in particular, in heterosexual dyads.

## Introduction

Behavioral coordination in social organisms, especially when responding to threats, is necessary for group survival, requiring the rapid integration of cues representing potential threats and social cues from conspecifics. Studies show that in animals and also in humans, social cues affect emotional states, which, in turn, alter the perception of these cues (Taylor 1981, Sandi and Haller 2015, Muroy, Long et al. 2016), suggesting an intertwining of the brain circuits responsible for processing affective and social information.

Several forms of socially modulated threat responses have been characterized in rodents (Morozov and Ito 2019). They include enhanced fear learning after exposure to stressed peers (Knapska, Mikosz et al. 2010), observational fear learning (Chen, Panksepp and Lahvis 2009, Jeon, Kim et al. 2010), fear conditioning by proxy (Bruchey, Jones and Monfils 2010, Jones, Riha et al. 2014), and fear buffering by non-fearful conspecifics (Kikusui, Winslow and Mori 2006, Kiyokawa, Takeuchi and Mori 2007, Guzman, Tronson et al. 2009). These forms, however, reflect delayed and unidirectional social effects, which are assessed mostly in a non-social context by testing single subjects individually or several days after the social interactions. In contrast, socially coordinated threat responses entail real-time, bidirectional cue exchange among animals responding to immediate threats simultaneously. This aspect of social behavior remains overlooked in current studies.

We developed a paradigm for studying coordinated threat responses in mice (Ito, Palmer and Morozov 2023), where fear-conditioned mouse dyads exposed to a continuous tone (120 s) as a conditioned stimulus (CS) show synchronized freezing patterns. The affective CS drives freezing in each mouse, but synchronization depends on social cues from the partner. This setup allows examination of how affective and social elements interact and shape behavioral synchronization. We observed more robust synchronization in male mice than females (Ito, Palmer and Morozov 2023), prompting further investigation into sex differences in fear synchrony. Additionally, we explored a newly identified coordinated defensive behavior, CS-induced affiliation.

Sex differences in behavioral coordination, as shown in various human studies (Weitz 1976, Lafrance and Ickes 1981, Tronick and Cohn 1989, Fujiwara, Kimura and Daibo 2019), likely reflect the distinct roles of males and females in handling social group threats (Van Vugt, De Cremer and Janssen 2007). Given the female’s primary role in nurturing and protecting progeny, Taylor and colleagues proposed that, unlike males, females evolved to “selectively affiliate in response to stress, which maximizes the likelihood that multiple group members will protect them and their offsprings” (Taylor, Klein et al. 2000). Given the well-documented sex differences in social and emotional processing (Taylor, Klein et al. 2000, Reyes, Valentino and Van Bockstaele 2008, Bangasser, Reyes et al. 2013, Bangasser and Cuarenta 2021), we hypothesized that the interplay between social and affective brain systems varies between sexes.

We examined this by altering affective states and social information in a fear synchrony paradigm. Dyads were stressed to change affective states, and familiar or unfamiliar mice pairings were used to vary social cues. The modulations resulted in sex-specific effects on fear synchrony and affiliative behaviors in same-sex dyads, implying sex differences in the interplay between the social and emotional systems. Interestingly, both synchrony and affiliation in opposite-sex dyads demonstrated resilience to stress and unfamiliarity, suggesting a unique and potentially adaptive mechanism that modulates the affective system in response to cues from the opposite sex.

## Results

### The social and non-social components of fear synchrony

We recently reported that mice in dyads synchronize conditioned freezing (Ito, Palmer and Morozov 2023); however, whether the synchronization relies exclusively on social cues has not been examined. Suppose two mice showed similar temporal freezing dynamics due to intrinsic freezing properties or any reason; our synchrony metric interprets these similar dynamics as a high degree of synchrony, even if it has nothing to do with exchanging social cues. Therefore, we investigated the non-social synchrony component through the following two approaches.

First, we compared synchrony between partners tested in the same chamber (PAIR) and in separate chambers isolated from each other without exchanging social cues (SINGLE). We fear-conditioned the mice and ran the two tests the following day. To mitigate the impact of repeated testing, we separated the two tests by a 5-hour interval and counterbalanced the testing sequences between “PAIR➜SINGLE” and “SINGLE➜PAIR” (**Fig1a**). The synchrony was significantly above 0 in PAIR (male: p<0.0001, female: p<0.001) but not SINGLE configuration in both sexes (**Fig1b**), suggesting a negligible non-social component.

**Fig 1.**
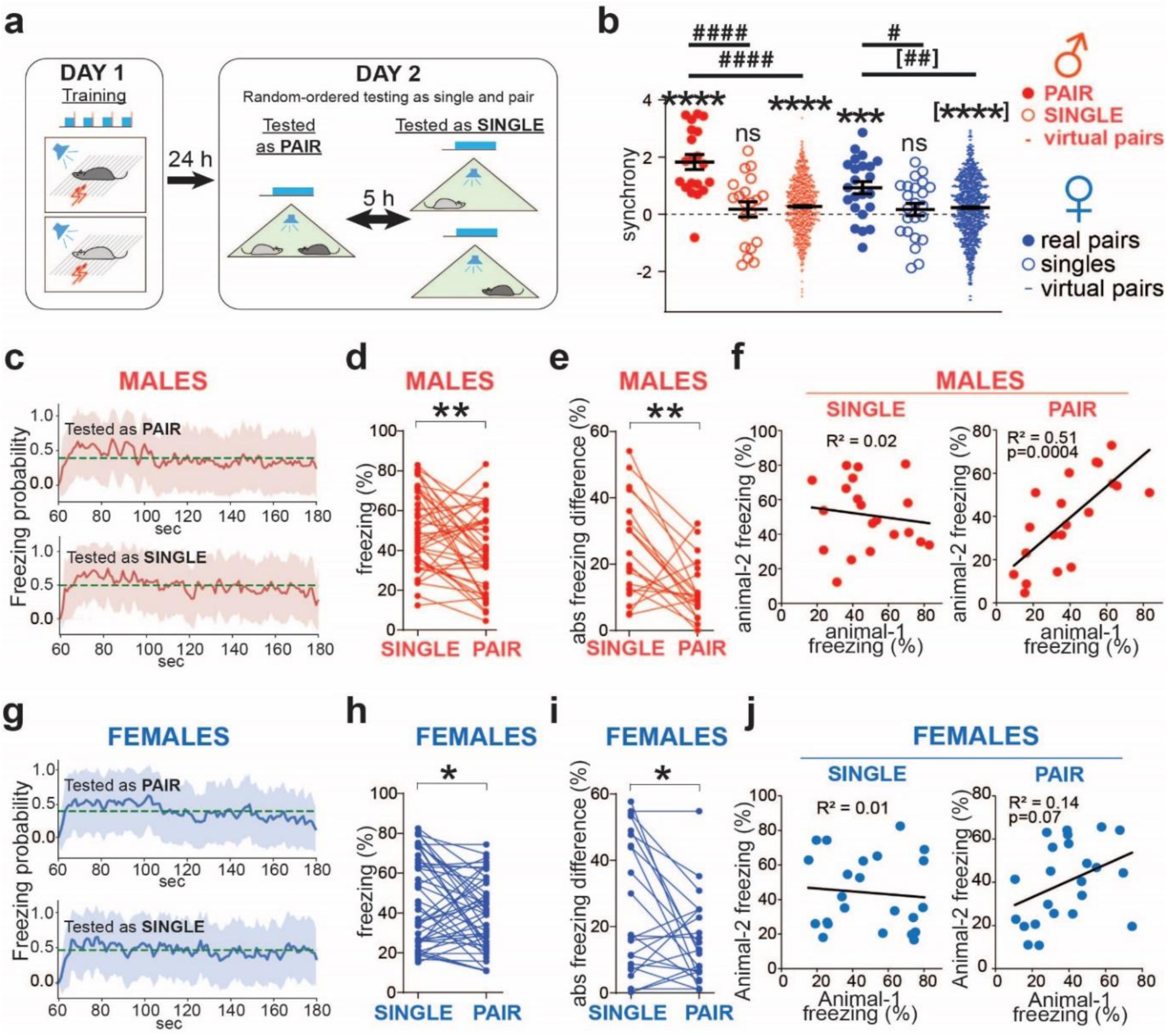
Male dyads coordinate freezing more strongly than females. **a** Scheme for testing the social modulation of freezing. Fear conditioning on Day 1, followed by counterbalanced sequential fear testing in either the ‘PAIR➜SINGLE’ or ‘SINGLE➜PAIR’ orders on Day 2. **b** Significant synchrony when tested together (PAIR) but not in isolation (SINGLE). However, the virtual pairs reveal a small non-social synchrony component. Horizontal bars indicate means±SEM. One-sample t-test comparing to 0: *** p<0.001, **** p<0.0001. Wilcoxon Signed Rank Test comparing to 0: [****] p<0.0001. Two sample t-test: ^#^ p<0.05, ^####^ p<0.0001. Mann-Whitney test: [^##^] p<0.01. Male dyads n=20, female dyads n=23, virtual male dyads n=760, virtual female dyads n=1012. **c, g** Averaged freezing probability during the CS period, computed using a sliding window of 1s. Dashed lines show the probability means. Shadowed areas represent SD (male n=40, female n=46). **d, h** Freezing levels of individual animals tested alone (SINGLE) or with a partner (PAIR). n=40 (males), 46 (females). Wilcoxon matched pairs test: * p<0.05, ** p<0.01. **e, i** Absolute freezing differences between partners tested individually (SINGLE) and together (PAIR). Male (n=20) and female (n=23) dyads. Wilcoxon matched pairs test: : * p<0.05, ** p<0.01. **f, j** Freezing levels of male partners correlated when tested together (PAIR) but not separately (SINGLE), and the correlations in females were not significant in both cases. The black lines show linear regression. The correlation coefficients (R^2^) and p-values are shown in the case of statistically significant or nearly significant correlations.

Nevertheless, it remained a concern that individually tested mice exhibited different freezing patterns from when tested with partners because of the isolation. To exclude such isolation effect, we computed synchrony in the virtual dyads – all possible combinations between mice tested as PAIR excluding real experimental pairs (20 male and 23 female real dyads yielded 760 male and 1012 female virtual dyads). It revealed small non-social synchrony components (**Fig1b**, virtual pairs, males: 0.27±0.03, females: 0.23±0.03, significantly positive due to overpowering by high N) and similar to the synchrony in the SINGLE configuration (males 0.17±0.27, females: 0.17±0.21). Both types of non-social synchrony were significantly smaller than the synchrony in the PAIR configuration. Meanwhile, computing the dynamics of the averaged freezing probabilities revealed a similar temporal pattern in both testing configurations: a rapid initial rise in about 5 seconds followed by a slow decline regardless of the partner’s presence or sex (**Fig1c, g**), presumably the cause of the identified small non-social synchrony components.

Finally, we examined whether mice synchronize to a non-social object – the Sphero Mini Robot Ball (Sphero, Hong Kong) (Solie, Contestabile et al. 2022), programmed to mimic precisely the freezing patterns from previously tested mice. The ball moves continuously during the pre-CS period and then stops and occasionally moves during CS, as the tested mice did (**Supplementary Fig1,** left). The mean synchrony with the ball was positive but not significantly different from 0 in both sexes (**Supplementary Fig1,** clean ball, males: 0.29±0.28; females: 0.42±0.22). To endow the ball with a social characteristic, we coated it with urine from a mouse of the opposite sex relative to the subject. Both sexes showed significant synchrony with the urine-pained ball (**Supplementary Fig1,** urinated ball, males: 0.79±0.30, p=0.024; females: 1.20±0.37, p=0.015).

**Supplementary Fig 1.**
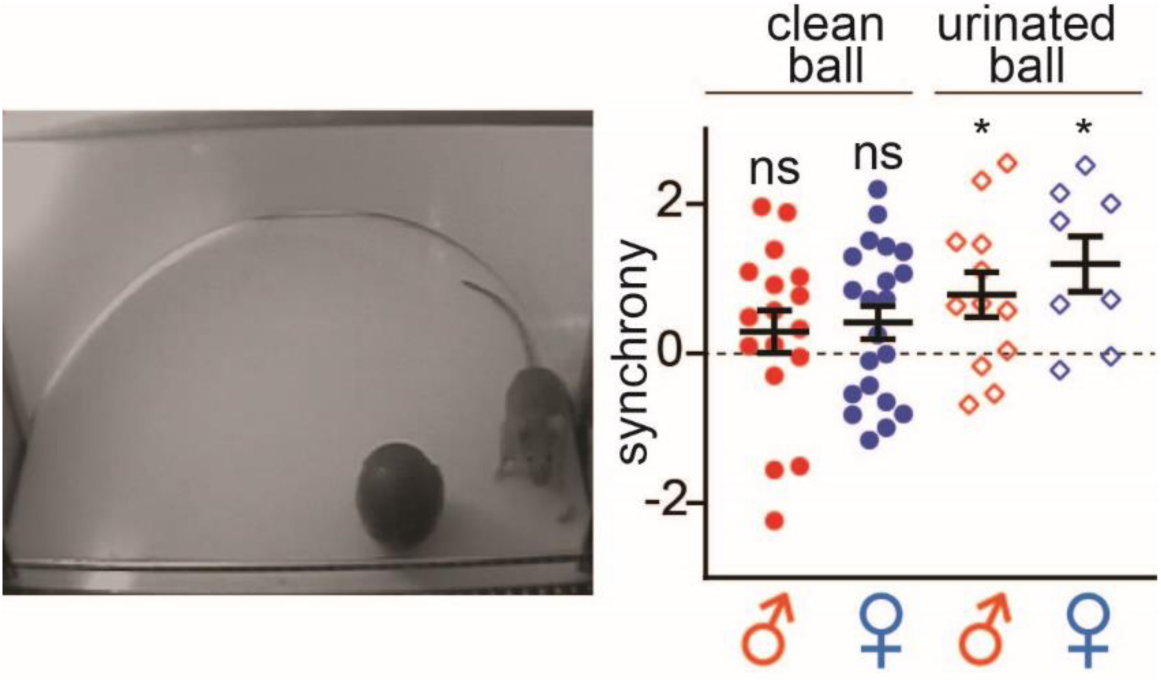
Mice do not synchronize with non-social ball, but urine-painting on the ball enables synchrony. **Left** Synchrony test with non-social ball. **Right** Summary of synchrony for each group (clean ball: n=17 (male), 21 (female), urinated ball: n=12 (male), 8 (female)). One-sample t-test comparing to 0: * p<0.05.

Those three results indicate that the observed synchronized freezing is driven primarily by social cues. Nevertheless, we updated our synchrony quantification by subtracting the non-social component computed from the virtual dyads in every experimental group.

### Equalization of fear response is more pronounced in males than in females

We further investigated how the conspecific presence influences the level of fear during fear retrieval. In line with previous studies (Kiyokawa, Takeuchi and Mori 2007, Fuzzo, Matsumoto et al. 2015), mice of both sexes demonstrated fear buffering by exhibiting reduced freezing in the presence of a partner, with this effect being more pronounced in males (p=0.007) than females (p=0.029) (**Fig1d, h**). Additionally, the absolute difference in freezing levels between partners in both sexes became smaller when tested as PAIR than when tested as SINGLE. Notably, this equalization effect was more pronounced in males (p=0.008) than in females (p=0.032), as shown in **Fig1e, i**. Furthermore, no correlations between partners’ freezing were observed when partners were tested in separate chambers (**Fig1f, j,** left SINGLE). In contrast, when tested together, the freezing levels of male partners showed a strong correlation (p=0.0004); however, this correlation was not significant in female pairs (p=0.07) (**Fig1f, j,** right PAIR).

These data indicate that social cues modulate fear levels towards equalization more strongly in male than female dyads.

### Brief stress diminishes fear synchrony in male dyads but augments it in female dyads

To induce a negative affective state, we subjected mice to 5-minute restraint stress one hour before the synchrony test (Lolait, Stewart et al. 2007, Newson, Pope et al. 2013) (**Fig2a**). For synchrony, the two-way ANOVA revealed a significant stress*sex interaction (F_(1,75)_ = 19.9, p<0.0001). The stressed males’ synchrony was lower than in unstressed males (p=0.0008) and did not differ from zero (**Fig2b left**). Unexpectedly, the stressed females’ synchrony was higher than in non-stressed females (p=0.0037) and well above zero (p<0.0001) (**Fig2b right**). For freezing, the two-way ANOVA revealed a significant stress*sex interaction (F_(1,154)_ = 11.3, p=0.001). Freezing levels did not differ between stressed and non-stressed males but were higher in stressed than non-stressed females (p<0.0001) (**Fig2c**). Merging data from stressed and unstressed dyads revealed a positive correlation between synchrony and freezing in female but not male dyads (**Supplementary Fig2**), consistent with the previous report (Ito, Palmer and Morozov 2023).

**Fig 2.**
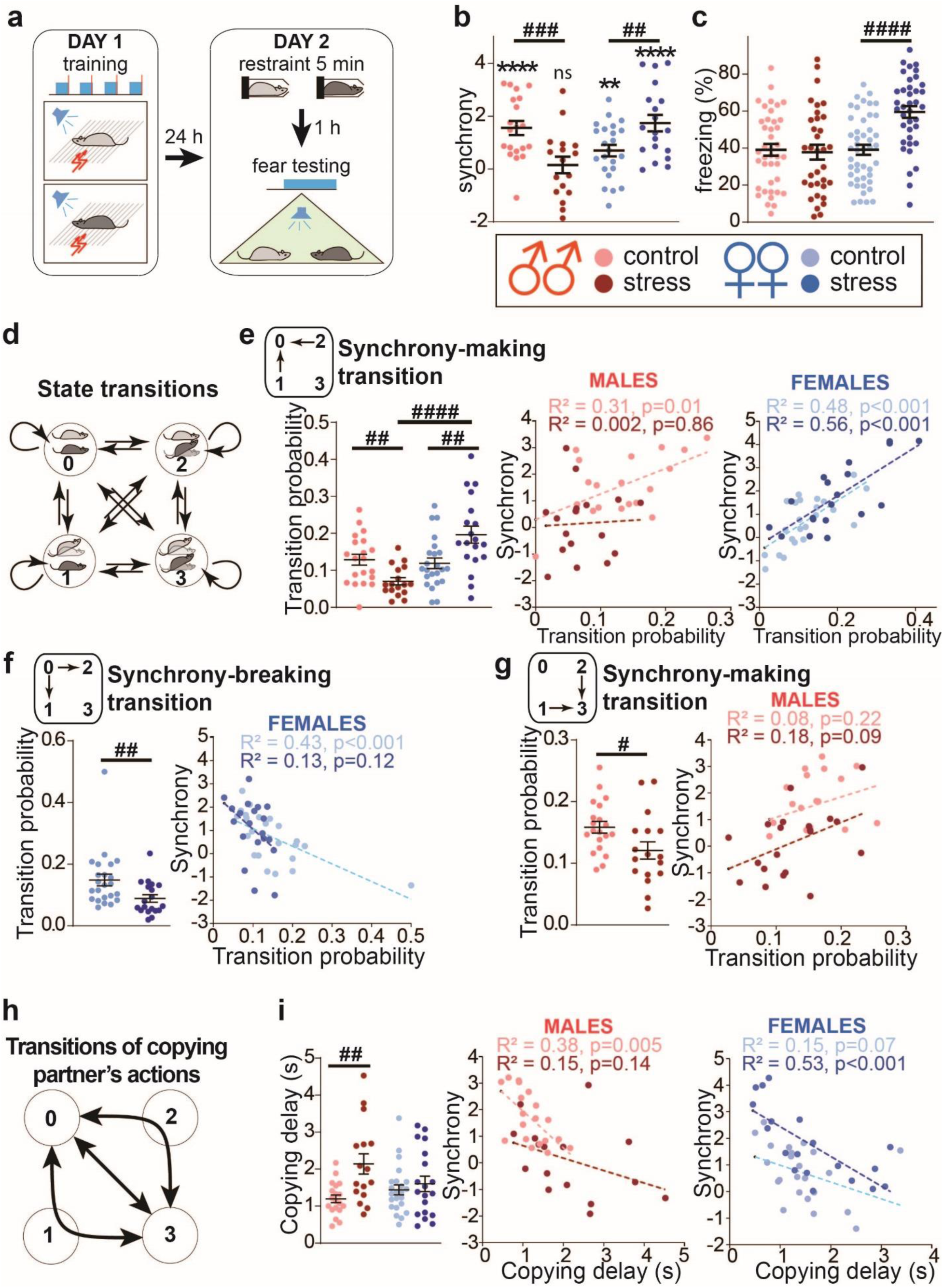
Sex-dimorphic effect of stress on fear synchrony and the underlying behaviors. **a** Scheme for testing stress effect on fear synchrony. Fear conditioning training on Day 1, and five min restraint followed by fear testing on Day 2. **b, c** Summary diagrams for synchrony (**b**) and average freezing (**c**). Colors representing sex (red: male dyads, blue: female dyads) and treatments (light symbols: control, males: n=20, females: n=23, dark symbols: stress, males: n=17, females: n=19). The color scheme is the same in the following panels. **d** Four behavioral states of a dyad (state 0,1,2,3): horizontal and angled mouse symbols represent freezing and moving mice, respectively. Arrows depict all possible transitions among the four states. **e-g** State transitions affected by stress as follows: 1/2➜0 (**e**), 0➜1/2 (**f**), and 1/2➜3 (**g**). **Left:** transition probability diagrams. **Right:** Transition probability vs. synchrony scatter plots and linear regressions. The Pearson correlation coefficients (R^2^) and significance values (p) are shown. **h** Diagram of copying behaviors as transitions between states 0 and 3 directly or via intermediate states 1 and 2. **i** Summary of stress effect on copying delay. **right:** Copying delays is defined as the duration of states 1 or 2. (control males: n=19, stressed males: n=16, control females: n=23, stressed females: n=19). **left:** Copying delay vs. synchrony scatter plots and linear regressions. The Pearson correlation coefficients (R^2^) and significance values (p) are shown. Horizontal bars on diagrams indicate means±SEM. **p<0.01, ****p<0.0001, one sample t-test compared to 0; ^#^ p<0.05^##^, p<0.01, ^####^ p<0.0001, two-sample t-test.

**Supplementary Fig 2.**
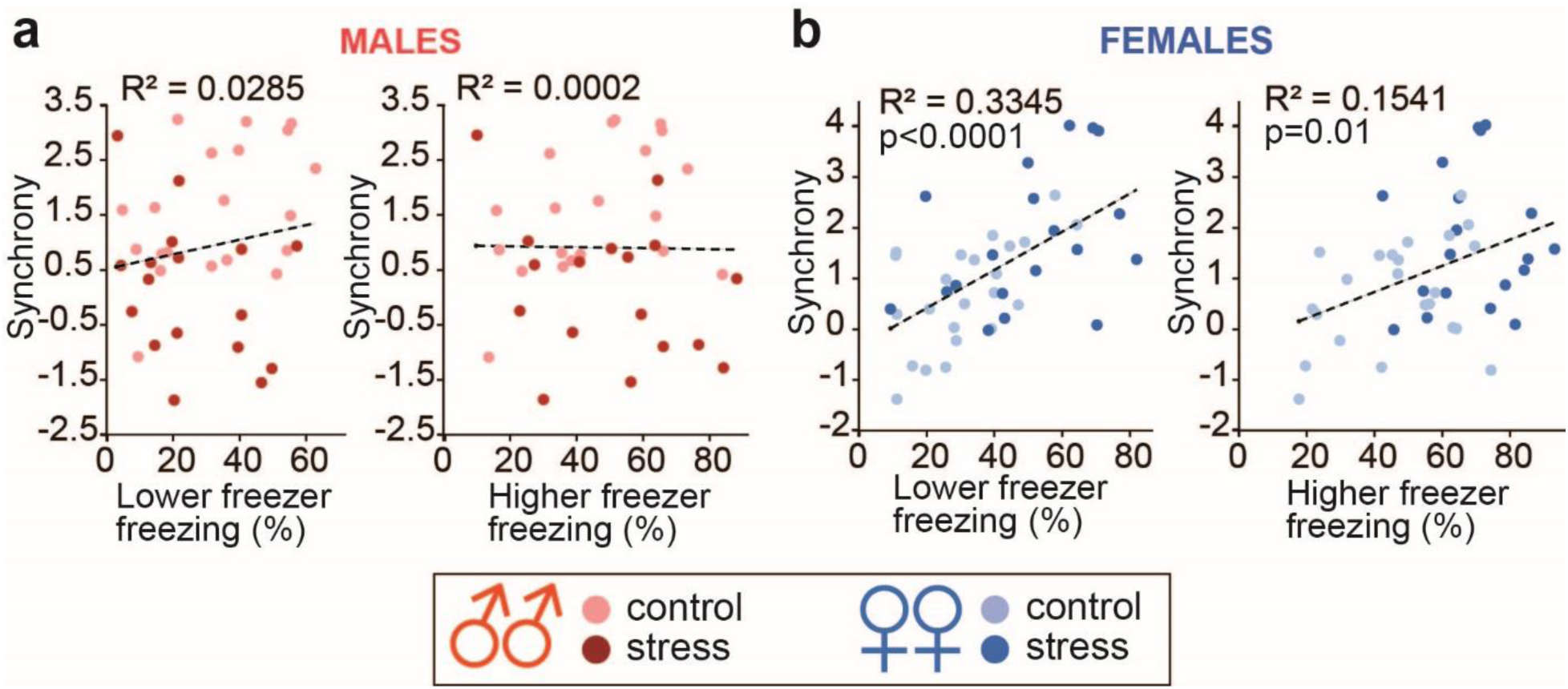
Freezing levels correlate with synchrony in females but not males. Scatter plot for synchrony vs. freezing levels from lower and higher freezing members. The color coding is the same in Fig1 (light symbols: control, males: n=20, females: n=23, dark symbols: stress, males: n=17, females: n=19). The regression lines, correlation coefficients (R^2^) and significant p-values are shown.

Meanwhile, the partners’ freezing levels no longer correlated in stressed males (**Supplementary Fig3a**), consistent with the synchrony loss. No such correlation was observed in females, regardless of stress. (**Supplementary Fig3b and Fig1i**) despite the high synchrony of stressed females. These findings suggest that brief stress weakens the social modulation of freezing in male dyads but strengthens it in females.

### Behavioral State Transitions and Delay Time for Copying Partner’s Actions that Account for Sex Differences in Fear Synchrony

To understand the behavioral process behind the stress effects on fear synchrony, we aimed to dissect the process into intricate behavioral states and their subsequent transitions. We used a Markov chain model, describing the dynamic of a dyad comprised of two animals - animal A and animal B - as transitions among four unique states (**Fig2d**): states 0, 1, 2, and 3. In state 0, animals A and B both freeze. In state 1, animal A freezes while animal B moves. Conversely, in state 2, animal A moves while animal B freezes. Finally, in state 3, both animals move. We designated states 0 and 3 as ‘congruent,’ symbolizing the same actions, while states 1 and 2 were labeled ‘incongruent,’ representing contrasting actions. We identified state transitions from the videos and computed each transition’s probability as the fraction of all transitions made from the corresponding initial states. Since the 1➜0 and 2➜0 transitions are behaviorally symmetrical, we computed a single probability value representing the combination of the two as 1/2➜0. The same was applied to the 1➜3 and 2➜3 transitions, represented as 1/2➜3. We then identified the stress-affected transitions and summarized them in **Fig2e-g**.

The probability of the synchrony-making transition 1/2➜0 was decreased by stress in males (p=0.0014) but increased in females (p=0.007), resulting in a significant difference between stressed males and females (p<0.0001) (**Fig2e left**), in line with the stress effects on synchrony. Furthermore, the transition probability correlated with synchrony in all groups, except the stressed males, in which stress skewed the probability towards the lower values (**Fig2e middle, right**).

Meanwhile, the probability of the synchrony-breaking transition 0➜1/2 was decreased by stress only in females (p=0.004, **Fig2f left**), which explains stress-enhanced synchrony in females. The probability negatively correlated with synchrony in control but not in stressed female dyads, similar to the 1/2➜0 synchrony-making transition of male dyads (**Fig2f right**). In males, stress did not significantly affect the 0➜1/2 transition, and the transition did not correlate with synchrony (**Supplementary Fig4a**).

Lastly, the probability of another synchrony-making transition 1/2➜3 was decreased by stress only in males (p=0.029), matched with the stress-abolished synchrony; however, that transition probability did not correlate with synchrony (**Fig2g**). Whereas, in females, the transition probability decrease did not reach significance, but interestingly, the probability correlated with synchrony in stressed dyads only (**Supplementary Fig4b**).

On top of the analyses above, we examined the delay in copying partners’ behavior as another behavioral measure that could explain stress effects on synchrony. Animals can synchronize by copying each other’s actions, for instance, initiating freezing after the partner begins to freeze. We formalized copying as the transitions between states 0 and 3, the two congruent states, directly or via non-congruent states 1 or 2 (**Fig2h**). Therefore, the duration in states 1 and 2 during those transitions represents the copying delay. The direct transitions 0**⇆**3 were assumed to represent the transitions that took less than the video sampling intervals of 250 ms (at four frames per sec), evaluated as the zero delay time.

Stress increased copying delay in male dyads (p=0.0024) but did not affect females (**Fig2i left**). The increased copying delay is consistent with the stressed male’s lower synchrony. Furthermore, the delay negatively correlated with synchrony in control but not in stressed male dyads (**Fig2i middle**), indicating that stress decouples synchronization from copying. In contrast, in females, the delay negatively correlated with synchrony in stressed but not in control dyads (**Fig2i right**), suggesting that stress makes synchrony dependent on copying. Notably, stress did not shorten the delay in female dyads, even though it enhanced fear synchrony. This discordance in female dyads implies that stress may affect their synchrony through a different mechanism from males.

These findings reveal stress effects on unique behavioral metrics in males and females, explaining sex-specific modulation of synchrony by stress.

### Conditioned Stimulus (CS) Promotes Social Affiliation, However, Stress Alters Male Behavior from Affiliation to Dispersion

Many social organisms exhibit coordinated social behavior, physically aggregating in response to perceived threats (Duranton and Gaunet 2016). However, rodents’ defensive aggregation or affiliation has only been reported in rats exposed to a predator’s odor (Bowen, Keats et al. 2012, Bowen, Kevin et al. 2013, Bowen and McGregor 2014), not in the popular Pavolovian fear conditioning. Therefore, we tracked the distance between the mice to determine whether CS onset facilitates proximity and whether the proximity between two animals influences fear synchronization. We chose the snout to represent the animal’s location, as its position and movement reflect the attentional orientation of the animal.

Accordingly, we compared the mean distances between two animals before and during the CS. **Fig3a** and **3d** depict the dynamics of the distances throughout the pre-CS and CS periods. First, during pre-CS, control (non-stressed) animals showed a larger distance in female dyads than in males (p=0.006). Stress increased the pre-CS distance in males (p=0.0002) but did not affect females. Then, CS decreased the distances in most groups (non-stressed males: p=0.012, non-stressed females: p=0.00067, stressed females: p=0.019); however, in stressed males, CS opposingly increased the distances (p=0.048) (**Fig3a, b, d, e**).

**Fig 3.**
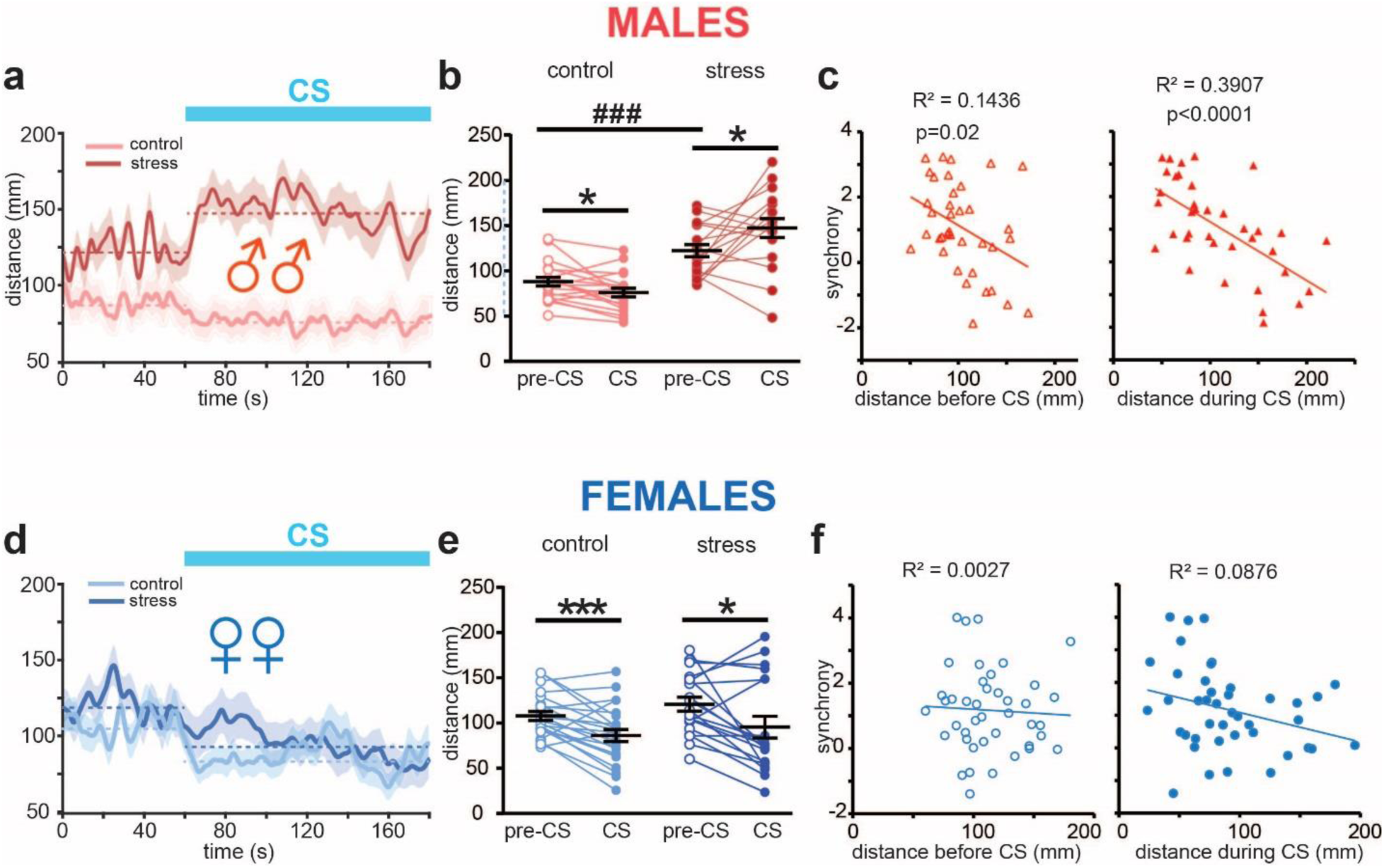
CS-induced affiliation predicts synchrony and is regulated by stress only in males. **a,d** Snout-to-snout mean distance dynamics along the testing session in males (**a**) and females (**d**), controls (light colors, males: n=20, females: n=23) or stressed (dark colors, males: n=17, females: n=19). The solid lines represent mean distances smoothed by Gaussian filter (the kernel SD = 5). The shaded areas depict SEMs. The borken lines indicate mean distances for the entire preCS and CS periods. **b, e** Mean distances during Pre-CS period and CS period in males (**b**) and females (**e**). **c,f** Scatter plots of synchrony vs. snout-to-snout distance before CS (**left**) and during CS (**right**) in merged stressed and non-stressed males (**c**) and females (**f**). Synchrony negatively correlated with distance only in males. The regression lines, correlation coefficients R^2^, and p-values for the Pearson correlation are shown in each scatter plot. *p<0.05, *** p<0.001, paired t-test; ^###^p<0.001, unpaired t-test; Means and SEM are shown by horizontal bars.

Next, a regression analysis on merged data from stressed and non-stressed dyads unveiled, only in male dyads, a negative correlation between fear synchrony and snout-to-snout distance (**Fig3c**), which was stronger during CS (R^2^=0.39, p<0.0001) than during pre-CS (R^2^=0.14, p=0.02). Such correlation was not observed in females (**Fig3f**).

Along the stress experiment, the CS-induced affiliation in Pavlovian fear conditioning, reported here for the first time, is revealed to have a male-specific link to fear synchrony.

### Unfamiliarity decreases fear synchrony in males and has sex-specific effects on freezing

Prior studies have shown that fear is modulated more effectively among familiar rodents (Jeon, Kim et al. 2010, Kiyokawa, Honda et al. 2014, Pisansky, Hanson et al. 2017). Additionally, it has been observed that familiar male mice exhibit a more robust transmission of fear compared to females (Pisansky, Hanson et al. 2017). Inspired by these insights, we examined the impact of familiarity on fear synchrony.

As predicted, unfamiliar male dyads displayed lower synchrony than familiar ones (p=0.04), but no differences were found in females; meanwhile, unfamiliar mice of both sexes exhibited significant fear synchrony (**Fig4a**). Despite decreasing synchrony in males, unfamiliarity did not significantly affect Markov state transition probabilities (data not shown); however, the copying delay was higher in unfamiliar than familiar male dyads (p=0.038) and negatively correlated with fear synchrony as well (**Fig4c**).

Unfamiliarity influenced freezing levels differently across sexes: In males, freezing levels were lower in unfamiliar dyads than familiar ones (p=0.027), while in females, the opposite effect was observed with higher freezing in unfamiliar than familiar dyads (p=0.01) (**Fig4b**). The two-way ANOVA detected a highly significant sex*unfamiliarity interaction (F(1,160)=10.5, p=0.0014). Despite the increase in the freezing levels of females, they did not correlate with synchrony (**Supplementary Fig5b**), unlike in familiar and stressed females (**Supplementary Fig2**). The decreased freezing levels of males did not correlate with synchrony either (**Supplementary Fig5a**). Furthermore, the freezing levels of unfamiliar partners did not correlate (**Supplementary Fig5c,d**), as in stressed animals in both sexes (**Supplementary Fig3**).

**Supplementary Fig 3.**
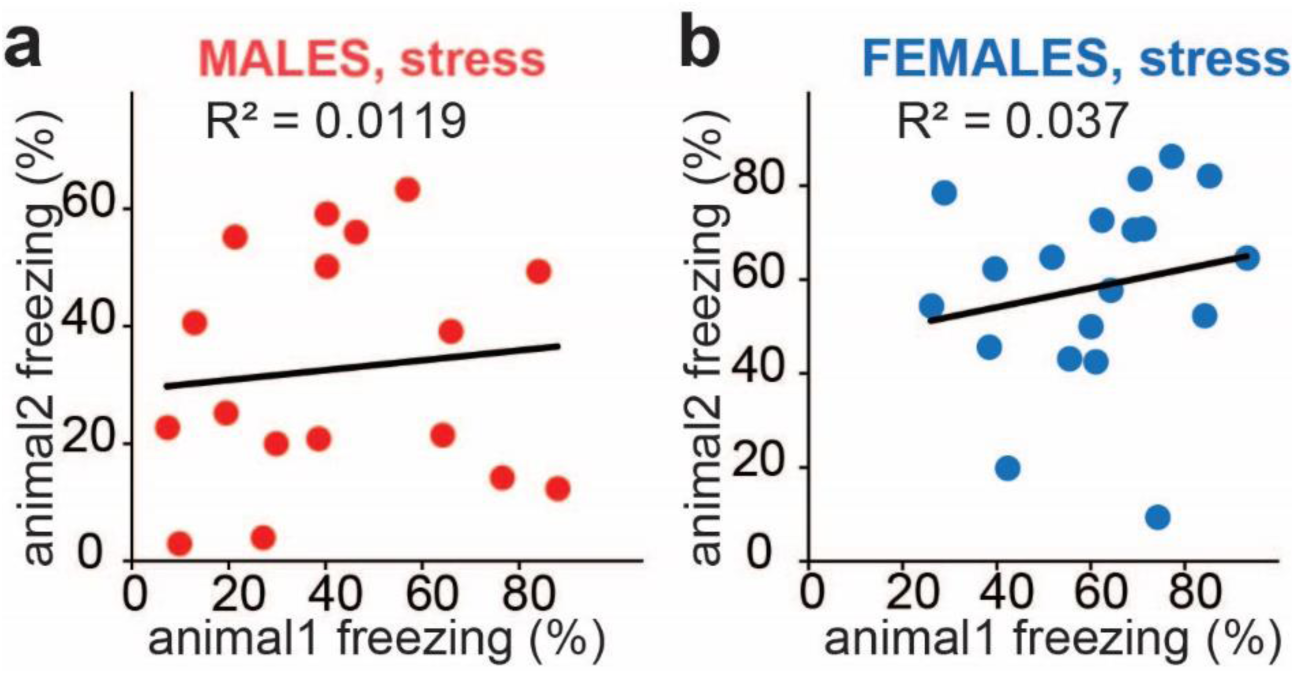
Freezing levels of stressed partners do not correlate. Scatter plots of partners’ freezing levels in male (**a**) (n=17) and female (**b**) (n=19) dyads. The regression lines and correlation coefficients (R^2^) are shown.

Meanwhile, another socially coordinated defensive behavior, CS-induced affiliation, was absent in both sexes (**Fig4d,e,g,h**). Intriguingly, the negative correlation between distance and synchrony emerged in unfamiliar females during the CS period, which was not observed in familiar and stressed females (**Fig4i**).

**Fig 4.**
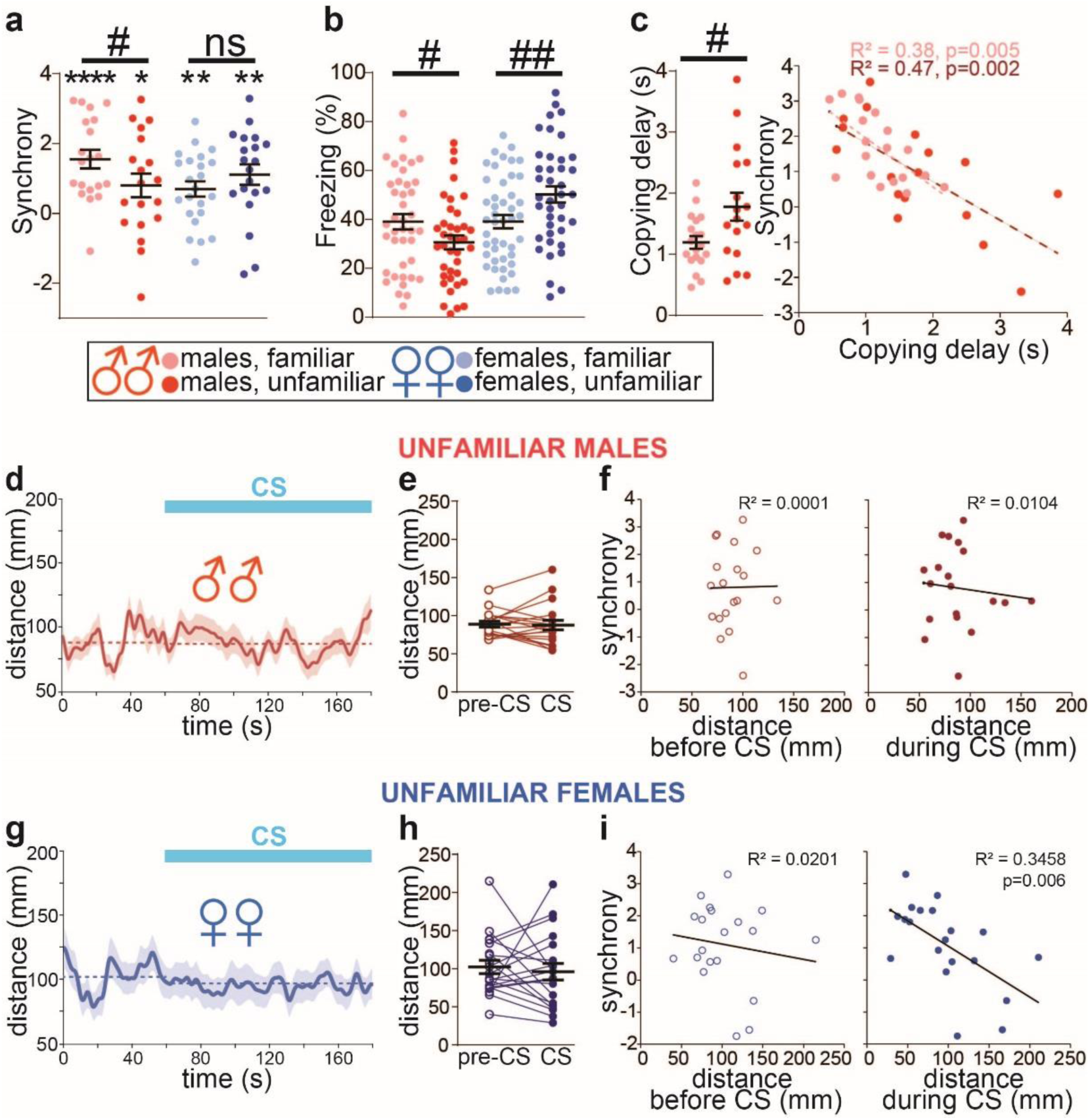
Unfamiliarity decreases fear synchrony in males and abolishes CS-induced affiliation in both sexes. **a** Synchrony in males (**left**) and females (**right**), familiar (light symbols, male dyads, n=20, female dyads, n=23) and unfamiliar (dark symbols, male dyads, n=19, female dyads, n=20). One sample t-test compared to 0: *p<0.05, **p<0.01, ****p<0.0001, two-sample t-test, one-tailed: ^#^p<0.05. **b** Freezing levels in males and females. Two-sample t-test: ^#^p<0.05, ^##^p<0.01. **c Left:** Copying delay in male dyads with unfamiliar partners (light red) is longer than with familiar ones (red). Two-sample t-test: ^#^p<0.05. **Right:** Synchrony negatively correlates with copying delay regardless of familiarity. The regression lines, correlation coefficients R^2^, and significant p-values for the Pearson correlation are shown. Means and SEM are shown by horizontal bars. **d-i** CS does not induce affiliation in unfamiliar animals. **d,g** Snout-to-snout mean distance dynamics along the testing session in males (**d**) (n=19) and females (**g**) (n=20). The solid lines represent mean distances smoothed by Gaussian filter (the kernel SD = 5). The shaded areas depict SEMs. The borken lines indicate mean distance during pre-CS and CS. **e,h** Mean distances before (preCS) and during (CS) CS in males (**e**) and females (**h**). **f,i** Scatter plots of synchrony vs. snout-to-snout distance before CS (**left)** and during CS (**right**). Synchrony negatively correlated with distance only in females during CS. The regression lines, correlation coefficients R^2^, and significant p-values for the Pearson correlation are shown. Means and SEM are shown by horizontal bars.

In line with published reports, unfamiliarity attenuated socially coordinated behaviors: CS-induced affiliation in both sexes and fear synchrony in male dyads. Meanwhile, unlike the stress effects, the unfamiliarity modulation of synchrony could not be explained using the decomposed behavioral metrics of the Markov transitions.

### No Impact from Stress or Unfamiliarity in Male-female Heterosexual Dyads

We finally examined the effects of stress and unfamiliarity on opposite-sex dyads. The “familiar” dyads comprised mice housed together for 5-7 days before training and testing. In contrast, the “unfamiliar” group consisted of mice housed with same-sex cagemates and tested with an unfamiliar partner of the opposite sex. For the “stress” group, the familiar dyads underwent a 5 min immobilization one hour before testing, as outlined in **Fig2a**.

Fear synchrony was highly significant across all three groups, with no discernable difference between groups (**Fig5a**). The mean freezing level was also consistent across groups (**Fig5b**). However, the partners’ freezing levels correlated only in the familiar dyads and degraded with stress and unfamiliarity (**Fig5c**). Similar to the unfamiliar same-sex dyads, the analysis of the Markov state transitions did not reveal any significant effect of stress or unfamiliarity (data not shown).

Another socially coordinated behavior, CS-induced affiliation, could not be detected by comparing the mean distances between partners (**Fig5d,e**). We also computed and compared the effect sizes (Cohen’s D) for the distance during pre-CS vs. CS periods to rule out a possibility that the highly fluctuating distance values obscured the CS-induced affiliation. The effect sizes were not significantly different from zero in all groups and were unaffected by stress or unfamiliarity (**Supplementary Fig6**). Meanwhile, the distance dynamics revealed gradual decreases during pre-CS periods (**Fig5d**) (mean distances in mm during the first vs. last five seconds of the pre-CS period, familiar: 86.0±1.4 vs. 62.7±1.4, p<0.0001; stressed: 102.7±2.7 vs. 68.1±1.6, p<0.0001; unfamiliar: 92.0±3.2 vs. 75.0±2.4, p<0.001). As a result, the average distances in heterosexual dyads were already shorter during the pre-CS period than in same-sex dyad groups. (Compared to unstressed males: p=0.07, unstressed females: p<0.0001, stressed males: p<0.0001, and stressed females: p<0.0001). Presumably, the affiliation between heterosexual partners occurring during pre-CS occluded any CS-induced affiliation.

Despite no apparent changes in the behavior metrics, merged data from the three groups revealed a moderate but significant negative correlation between synchrony and snout-to-snout distance during CS but not preCS (**Fig5g**). This suggests that proximity enhances synchrony in heterosexual dyads as well.

## Discussion

This study uncovered differences between sexes in how stress affects socially coordinated responses to conditioned threats. It provided behavioral evidence that affective and social brain systems interacted differently in males and females. Intriguingly, male-female heterosexual dyads displayed resistance to stress modulation akin to that observed in female same-sex dyads, suggesting that the opposite-sex cues negated the stress impact on the male brain.

We examined socio-emotional interactions by investigating two socially coordinated defensive behaviors that emerged when individually fear-conditioned mice were tested as dyads responding to CS. The first behavior was the synchronization of freezing bouts (Ito, Palmer and Morozov 2023), and the second one was a newly identified CS-induced affiliation.

We made five notable findings by manipulating social cues and the animals’ emotional states. 1) In same-sex dyads with familiar partners, males synchronized freezing bouts more strongly than females. 2) Prior brief immobilization stress abolished synchrony in males but enhanced it in females. 3) Stress switched male but not female dyads from CS-induced affiliation to dispersion. 4) Freezing synchrony correlated negatively with the distance between male but not female partners. 5) Male-female heterosexual dyads were resilient to stress and maintained high freezing synchrony and affiliation. In addition, only in males did unfamiliarity attenuate synchrony.

### Mice coordinate freezing levels more strongly in male than female dyads

Our study extends the recent findings of weaker females’ freezing synchrony as the coordination of freezing patterns (Ito, Palmer and Morozov 2023) by examining the coordination of freezing levels across three key comparisons. Firstly, we observed lower freezing levels in both sexes when tested in dyads versus individually (**Fig1d,h**), suggesting social buffering. Secondly, we noted a decrease in the variance between partners’ freezing levels (**Fig1e,i**), indicating freezing equalization. Both equalization and buffering were more significant in males than females. Thirdly, a significant correlation in freezing levels was seen in male dyads but not solo tests (**Fig1f**). In contrast, this correlation was negligible in females, reflecting less effective fear buffering and freezing equalization, thereby indicating weaker coordination of freezing levels compared to males.

### Sex-specific effects of stress on socially coordinated defensive behaviors

We applied acute stress to test how the emotional state modulates the socially coordinated freezing. In males, stress diminished both the coordination of freezing patterns (synchrony) and freezing levels (**Fig2b, Supplementary Fig3**). Conversely, stress enhanced freezing pattern coordination in females, yet it didn’t facilitate freezing level coordination, as evidenced by the lack of correlation in partners’ freezing. These results demonstrate that stress affects males’ and females’ abilities to assimilate social and emotional cues differently and imply that separate brain circuits govern the coordination of freezing levels and patterns.

Analyzing dyads’ behavioral state transitions helps understand the behavioral process underlying the sex-specific stress effects. The most notable and sex-dimorphic effect was on the “synchrony-making” 1/2➜0 transition, in which the non-freezing animal starts freezing alongside a freezing partner. Stress decreased that transition in males but increased it in females (**Fig2e**).

Effects of stress on other behavioral transitions were found only in one of the sexes. Thus, stress decreased the “synchrony-breaking” 0➜1/2 transition in females (**Fig2f**) but had no effect in males, where the transition did not correlate with synchrony (**Supplementary Fig4a**). In addition, stress decreased the 1/2➜3 transition in males but not females. Since the 1/2➜3 transition did not correlate with the male’s synchrony (**Fig2g**), its modulation does not explain the male-specific synchrony loss. Lastly, stress increased copying delay in males, consistent with the diminished synchrony, while in females, despite an increase in synchrony, there was no change in copying delay (**Fig2i**). These findings indicate that the observed sex differences in stress-induced fear synchrony stem from three key behavioral modifications: 1) the symmetrically contrasting impacts on the 1/2➜0 synchrony-making transition, 2) a female-specific reduction in the 0➜1/2 synchrony-breaking transition, and 3) male-specific increases in copying delay. Future studies involving neuronal activity recordings could be instrumental in pinpointing the neural circuits responsible for these specific behavioral dynamics.

**Supplementary Fig 4.**
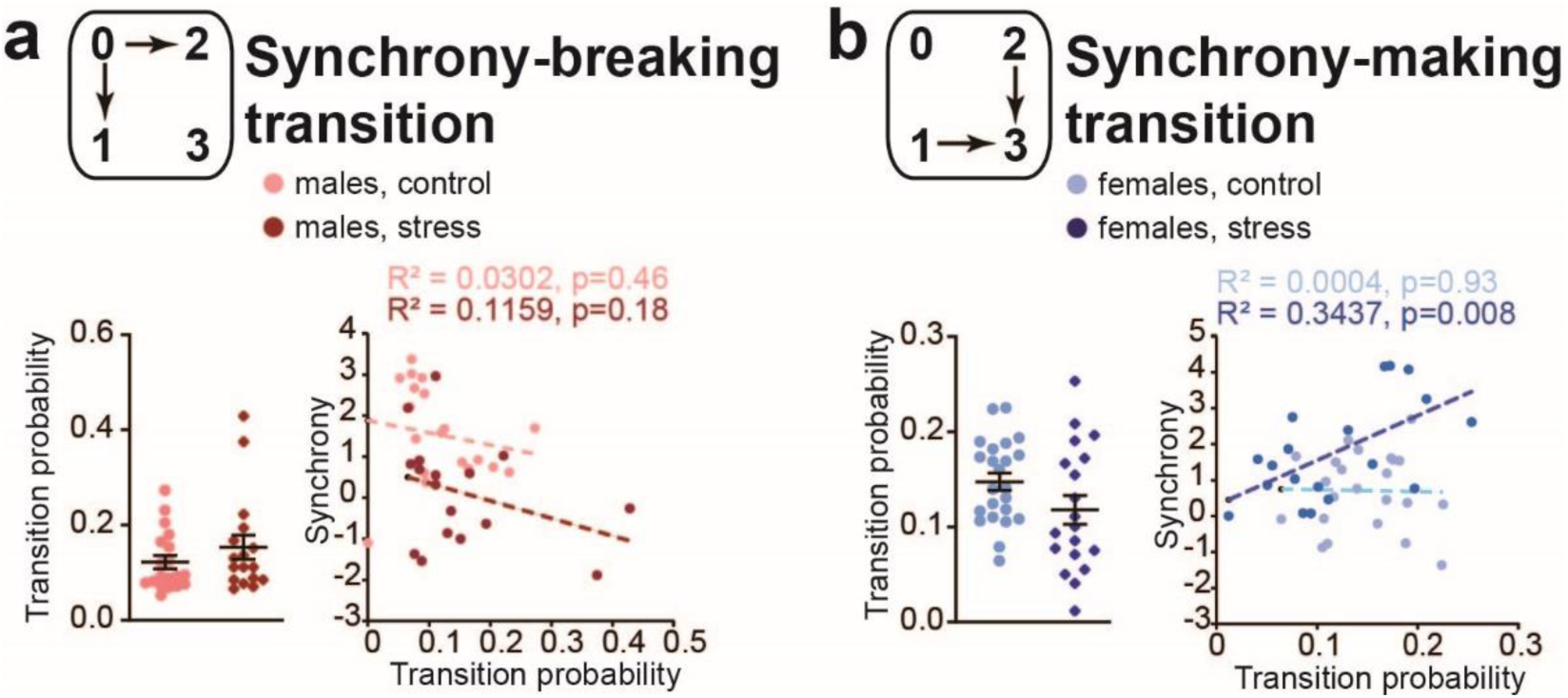
State transitions insensitive to stress in one of the sexes. **a** Data for the 0➜1/2 transition in males. **Left:** Transition probability in control (light color) and stressed (dark color) dyads. **Right:** Transition probability vs. synchrony scatter plots and linear regression lines. The Pearson correlation coefficients (R^2^) and significance values (p) are shown. **b** Data for the 1/2➜3 transition in females are shown as in **a**.

### CS-induced affiliation and its modulation by stress

We analyzed the distance between partners, hypothesizing that closer proximity enhances the exchange of social cues and increases synchrony. We found that both male and female mice exhibited CS-induced affiliation, demonstrated by reduced distances between partners (**Fig3a,d**), marking the first evidence of such behavior in mice. This phenomenon is akin to defensive aggregation seen in wild species (Duranton and Gaunet 2016), stress-induced affiliation in humans (Taylor, Klein et al. 2000), and predator odor-induced aggregation in rats (Bowen, Keats et al. 2012, Bowen, Kevin et al. 2013, Bowen and McGregor 2014). As predicted, synchrony in males correlated negatively with distance between partners, and the correlation was stronger during the CS than during the non-threatening pre-CS period (**Fig3c**). Surprisingly, this correlation was absent in females (**Fig3f**), suggesting that the exchange of social cues in females might be less reliant on physical proximity within the confines of our test chamber.

Stress interestingly influenced affiliation behaviors depending on CS. Stress reduced affiliation in both sexes during the non-threatening preCS period. This aligns with previous studies showing diminished prosocial behaviors in rodents under stress, particularly in unfamiliar environments (Sandi and Haller 2015), such as reduced social approach (Zain, Pandy et al. 2019) and empathy for pain (Zain, Pandy et al. 2019), though Muroy et al. (Muroy, Long et al. 2016) observed stress-enhanced affiliation in a familiar setting. Contrarily, stress caused males to shift from affiliation to separation during CS exposure, but this effect was not observed in females (**Fig3a,b,d,e**).

We are not aware of reports of sex differences in threat-induced affiliation in rodents; however, human studies revealed varying affiliative behaviors depending on the degree of emotional arousal, anxiety traits and gender. For instance, individuals with high trait anxiety tend to shun affiliation when highly aroused but seek it when less aroused. Conversely, those with low trait anxiety behave opposingly (Teichman 1974). Furthermore, stress appears to influence affiliation in a gender-specific manner, enhancing it in women while reducing it in men (Zucker, Manosevitz and Lanyon 1968, Bull, Burbage et al. 1972, Taylor, Klein et al. 2000, Sherman, Rice et al. 2017). These observations led Taylor and colleagues to suggest that the typical male response to stress is a fight-or-flight reaction. In contrast, females increase affiliative behaviors (Taylor, Klein et al. 2000). Our findings in mice seem to support Taylor’s hypothesis, suggesting similar gender-specific stress responses across species.

It seems likely that affiliation impacts freezing synchrony. Indeed, the CS-induced distancing observed in stressed males aligned with impaired fear synchrony, and there was a notable negative correlation between synchrony and distance. This implies that stress-induced social avoidance may reduce freezing synchrony in males. However, stress enhanced synchrony in females (**Fig2b**) without promoting closer proximity (**Fig3e**), and no correlation was found between synchrony and distance. This indicates that factors beyond mere physical proximity, possibly social arousal, as discussed in the following paragraph, play a pivotal role in determining females’ synchrony.

### Potential mechanisms of the sex-specific stress effects

The exact mechanisms behind the sexually dimorphic effects of stress are not fully understood, but several stress-responsive molecules (Bangasser and Cuarenta 2021) could play a role. For instance, oxytocin, which is released in response to stress (Bangasser and Cuarenta 2021), promotes prosocial behaviors in females, possibly due to estrogen enhancing oxytocin functions (McCarthy 1995). In contrast, oxytocin is linked to anxiety-like behaviors in males (Li, Nakajima et al. 2016) and is known to increase aggregation in male rats in response to predator odors (Bowen and McGregor 2014), suggesting its role in CS-induced aggregation and the sex-specific effects of stress.

Another key factor is the CRF-1 receptor in the locus coeruleus, which shows sex-dependent regulation: it desensitizes in males but remains active in females (Bangasser, Reyes et al. 2013). This sustained activation in females might increase social arousal, facilitating fear synchrony.

At the circuit level, stress decreases neuronal activity in the CA1, CA2, and CA3 areas of the hippocampus of males but not females (Lin, Ter Horst et al. 2009), potentially explaining the male-specific decrease in affiliation and synchrony, especially since freezing synchrony relies on the ventral hippocampus (vHPC) (Ito, Palmer and Morozov 2023). Unraveling these sex-specific molecular pathways that bridge social and emotional circuits is essential for a deeper understanding of sex differences in fear synchronization.

### Moderate effects of unfamiliarity

Unlike the pronounced effects of stress, the unfamiliarity of partners moderately reduced synchrony and only in male dyads. In these pairs, unfamiliarity also disrupted the correlation between partners’ freezing levels and led to an increase in copying delay. However, it did not significantly affect Markov transitions (**Fig4a,c and Supplementary Fig5c,d**). This result is consistent with previous research on observational fear learning, fear conditioning by proxy, and fear buffering, which showed that social modulation of threat responses is less effective with unfamiliar conspecifics (Jeon, Kim et al. 2010, Kiyokawa, Honda et al. 2014, Pisansky, Hanson et al. 2017, Agee, Jones and Monfils 2019).

In contrast, the unfamiliarity of partners did not affect female dyads, aligning with prior findings of unaffected observational fear learning in unfamiliar female subjects (Pisansky, Hanson et al. 2017). The sex-specific effect of unfamiliarity can arise from differences in perception of same-sex conspecifics. Male mice living in harems in the wild view unfamiliar males as threats. Such perception likely interferes with freezing coordination. In contrast, female mice, known for minimal intrasexual competition (Pandolfi, Scaia and Fernandez 2021), may pay less attention to same-sex conspecifics, leading to a negligible effect of unfamiliarity but lower than in male synchrony. While Blanchard’s lab noted lower sociability in females (Defensor, Pearson et al. 2011), indicating reduced social attention, other studies (Yang, Clarke and Crawley 2009, Silverman, Tolu et al. 2010) report no significant differences in sociability between sexes, suggesting the issue is more nuanced.

### The resilience of heterosexual dyads

The male-female dyads displayed relatively high levels of fear synchrony. These partners also maintained close proximity during the basal pre-CS period, which left little room for further reduction in distance by CS. Notably, neither stress nor familiarity impacted their synchrony or affiliation (**Fig5**). This resilience of coordinated threat responses and affiliation in opposite-sex dyads can be advantageous for offspring production and species survival, akin to the female-female affiliation in Taylor’s theory (Taylor, Klein et al. 2000).

**Fig 5.**
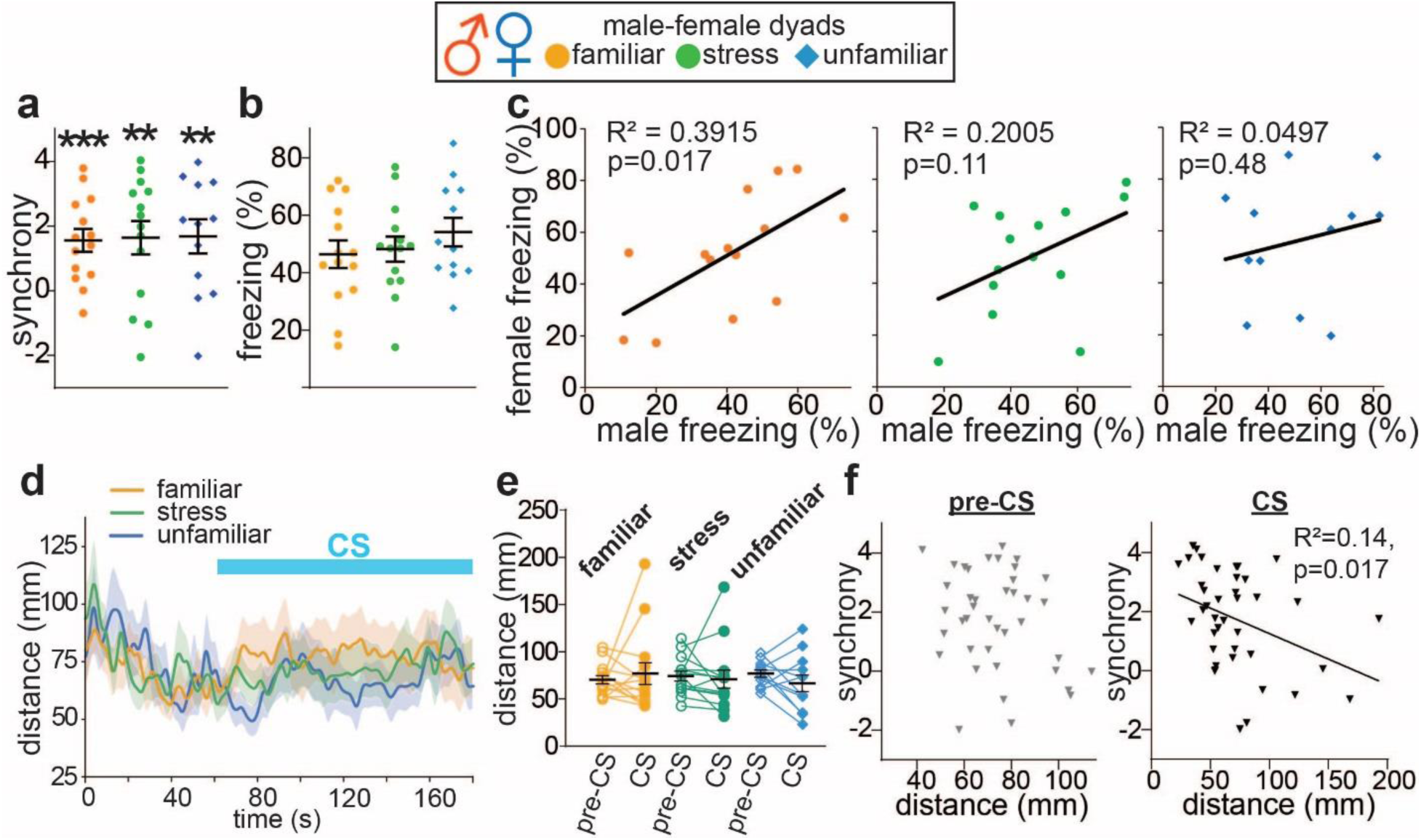
Stress and familiarity do not affects behaviors of male-female dyads. **a** Synchrony in male-female dyads composed of familiar (orange, n=14), stressed (green, n=14), and unfamiliar animals (blue, n=12). The color coding of groups is the same in all panels. **b** Average freezing. **c** Scatter plots of the female freezing as the function of the male freezing show correlation only between familiar unstressed partners. **d** Snout-to-snout mean distance dynamics during the testing session as in **Fig3**. **e** Snout-to-snout mean distances before CS (preCS) and during CS (CS). **f** Scatter plots of synchrony as the function of distance between partners before CS (**left**) and during CS (**right**) (n=40). ** p<0.01, ***p<0.001, one sample t-test compared to 0. Means and SEM are shown by horizontal bars.

**Supplementary Fig 5.**
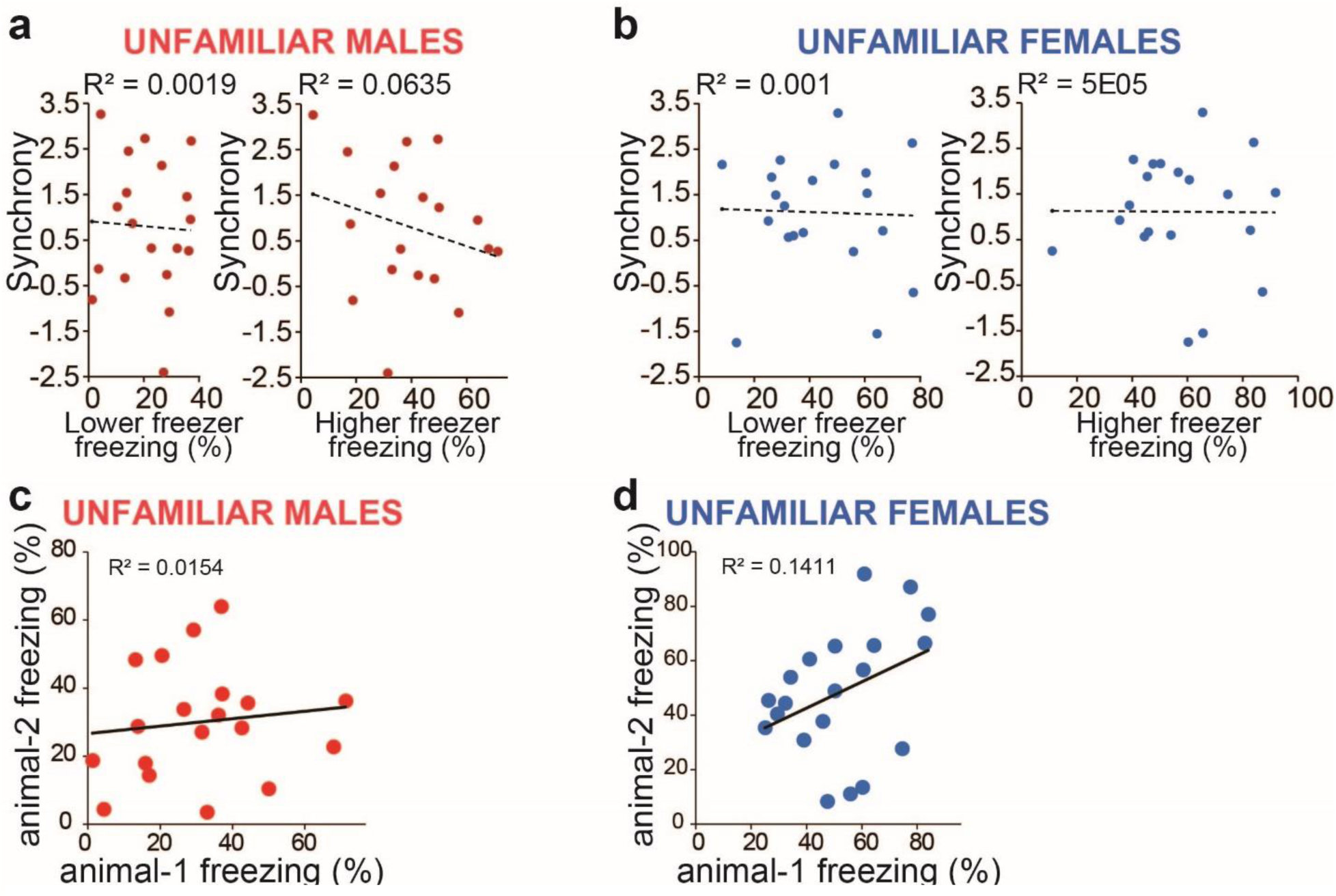
Unfamiliar dyads did not show correlations in freezing properties or CS-induced affiliation. **a, b** Freezing levels do not correlate with synchrony in both sexes. Scatter plot for synchrony vs. freezing levels from lower and higher freezing members of unfamiliar male dyads (**a**) and unfamiliar female dyads (**b**) (male dyads, n=19, female dyads, n=20). **c,d** Lack of correlation between freezing levels of unfamiliar partners in males (**c**) and females (**d**). The regression lines and correlation coefficients R_2_ are shown in each scatter plot.

**Supplementary Fig 6.**
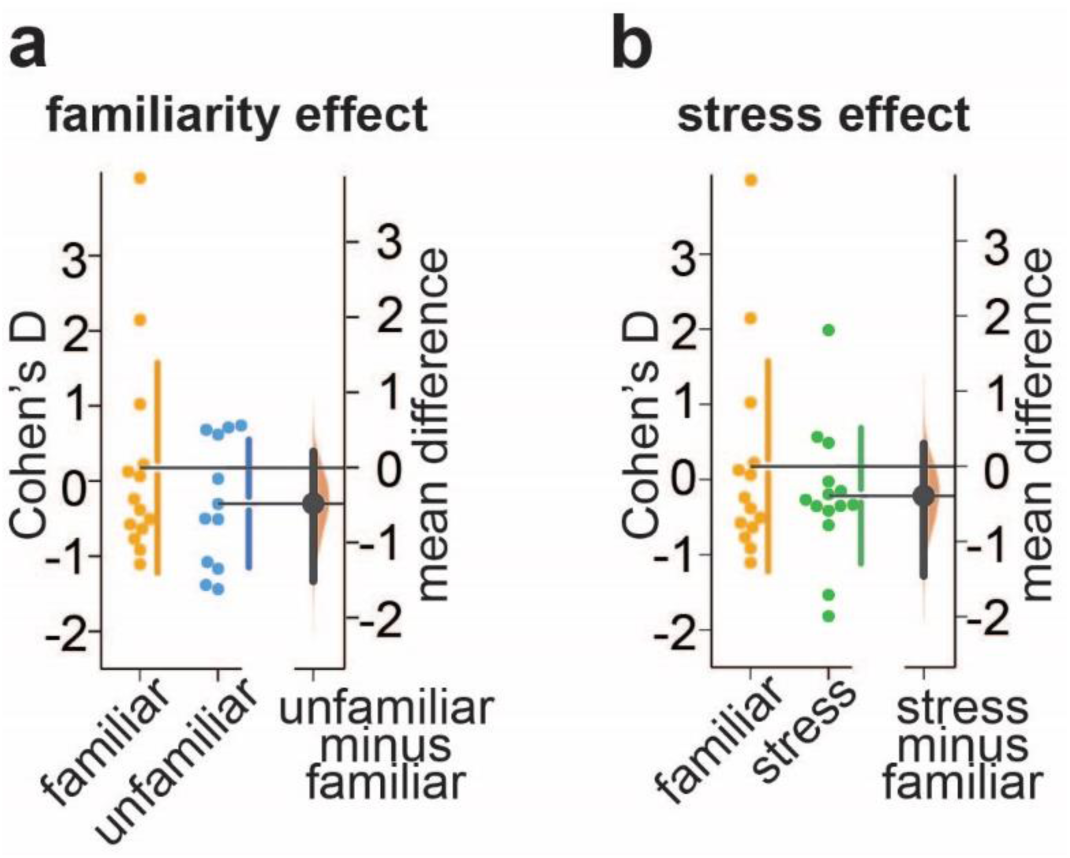
No significant CS-induced affiliatiion in heterosex dyads. **a,b** The effects from familiarity (**a**) and stress (**b**) on the effects sizes (Cohen’s D) for distances compared between pre-CS and CS for each dyad. In each panel, **Left:** Cohen’s D effect sizes for each distance. **Right:** the mean differences of Cohen’s D. Bootstrap 95% confidence intervals (vertical lines) and the resampled distributions of the mean difference (orange shadow) computed by DABEST (Ho, Tumkaya et al. 2019). The mean difference of 0 is aligned with the familiars’ Cohen’s D means.

Remarkably, the impact of heterosexual pairing appears more pronounced on the male brain, wherein female cues seem to counteract the disruptive effects of stress and unfamiliarity on males’ ability to affiliate and synchronize with a partner. This resilience could arise from robust social attention towards the opposite-sex partner and motivation to approach.

The exploration of the underlying circuit mechanism presents an intriguing avenue for research. Opposite-sex chemosensory signals are primarily transmitted via the vomeronasal pathway, activating sex-encoding neurons in the medial amygdala, hypothalamus, and prefrontal cortex (Li and Dulac 2018, Kingsbury, Huang et al. 2020). These signals eventually reach the circuits underlying coordinated defensive behaviors, such as the anterior hypothalamus and medial preoptic area, which can drive affiliation (Fukumitsu, Kaneko et al. 2022, Jabarin, Dagash et al. 2023), as well as the ventral hippocampus (vHPC) and basal amygdala (BA), which mediate freezing synchrony (Ito, Palmer and Morozov 2023). Understanding these circuit mechanisms will provide valuable insights into the neural basis of the opposite-sex effects on coordinated defensive behaviors and differences between same-sex male and female dyads. Furthermore, it can uncover sex-specific modulation of social behaviors and socio-emotional integrator by adverse experiences relevant to psychiatric disease.

## Materials and methods

All experiments were performed according to a Virginia Tech IACUC-approved protocol.

### Animals

Breeding trios of one C57BL/6N male and two 129SvEv females produced 129SvEv/C57BL/6N F1 hybrid male and female mice weaned at p21 and housed four littermates per cage of the same sex as described (Ito, Erisir and Morozov 2015). Subjects underwent tests at p75-p90. The same-sex familiar dyads were formed from the littermates 7 days before training and placed in cages with fresh bedding, which was not changed until the completion of testing. For the same-sex unfamiliar dyads, the mice were housed in pairs of littermates for 7 days before testing but paired randomly with unfamiliar non-littermates from different cages during testing. The heterosexual familiar dyads comprised mice housed together for 5-7 days before training and testing from the pool of the virgin animals, initially housed in groups of 4 same-sex littermates per cage. The unfamiliar heterosexual dyads consisted of mice always housed in groups of same-sex littermates.

### Fear conditioning

Mice in each dyad were trained independently in two separate conditioning chambers (Med Associates, St. Albans, VT) and then tested together as dyads in a single chamber as described (Ito, Palmer and Morozov 2023). For training, each animal received four pairings of the conditioned stimulus (CS: a 30 s, 8 kHz, 80 dB tone) and unconditioned stimulus (US: a 0.5 mA 0.5 s electrical shock co-terminated with CS) given in variable intervals (60-180 s). Cued fear was tested one or two days later in a new context. The animals spent the first 1 min without CS and then 2 min with CS. Videos were recorded at four frames per second, exported as AVI files with MJPEG compression using the Freezeframe system (Actimetrics, Wilmette, IL), and then converted to the mp4 format using a Python script.

### Immobilization stress

Mice were placed inside the 50 ml Falcon tubes with the 8 mm opening at the bottom to allow breathing. The animals were returned to the home cage 5 min later.

### Quantification of freezing, freezing synchrony and partners’ freezing correlations

Freezing was defined as the absence of all observable movement of the skeleton and vibrissae, except for those related to respiration (Fanselow 1980) during at least four consecutive video frames (1s). Freezing synchrony was defined as the standardized difference between the observed and chance freezing overlaps and quantified as previously described (Ito, Palmer and Morozov 2023). Briefly, annotators, unaware of the treatment of the animals, manually recorded the first and last video frames of each freezing bout using a Python script. From the annotation, another Python script calculated the freezing duration for each animal and then freezing overlap, chance overlap, and freezing synchrony for each dyad. To test the correlation between partners’ freezing levels, we assigned each dyad member to the two groups, animal 1 and animal 2, so that both groups had similar mean and distribution of freezing levels to avoid spurious correlation due to unbalanced assignment. We first sorted the dyads by their mean freezing levels, second, identified the higher and lower freezer animals in each dyad, and third, performed alternated assignment of the high or low freezers as animal 1 or 2 along the dyad’s freezing mean. The resulting animal 1 and animal 2 groups were balanced by freezing means and numbers of high and low freezers.

### Quantification of transitions among behavioral states

The behavioral states were sampled with every video frame (4 frames per second), giving 480 samples during 2 min of the CS period. The transition probabilities were calculated by dividing the counts of the transition of interest by the total number of all transitions from the corresponding initial state, including the self-loop back transitions to the same state. Since the 1➜0 and 2➜0 transitions are behaviorally symmetrical, we computed a single probability value summing the two probabilities as 1/2➜0. The same was applied to the pairs of 0➜1 and 0➜2 transitions, the 1➜3 and 2➜3 transitions, and the 3➜1 and 3➜2 transitions, represented as 0➜1/2, 1/2➜3 and 3➜1/2, respectively.

### Animal tracking and distance estimation

For each video frame, the X-Y coordinates of the snout projecting on the chamber floor were estimated using a triangular similarity-based formula, using the pixel coordinates of the snout, the three-chamber corners and the 3D coordinates for the corners. The distance between the snout projections was calculated using the X-Y coordinates.

### Data analysis

Statistical analyses were performed using GraphPad Prism 10 (GraphPad Software, La Jolla, CA). Normality was tested using the Shapiro–Wilk test. Datasets with normal distribution were compared using the one-sample or two-sample t-test. The datasets with non-normal distribution were compared using the Wilcoxon signed-rank test and Mann– Whitney test. All the tests were two-sided except when mentioned otherwise. The two-tailed p-value was calculated for the Spearman correlation analysis. The effects were deemed significant with p < 0.05.

## Data and code availability

All primary data, including video files, are available from authors upon reasonable request. In addition, codes for data analysis and statistics will be provided with example data as part of the replication package after publication.

## Acknowledgments

The study was supported by an NIH grant R01MH120290 to AM.

## Notes

### Competing Interest Statement

The authors have declared no competing interest.

### Summary of Updates

The Acknowledgments section with NIH funding information was added as follows: "The study was supported by an NIH grant R01MH120290 to AM."

